# Chronically radiation-exposed survivor glioblastoma cells display poor response to Chk1 inhibition

**DOI:** 10.1101/2022.04.16.488495

**Authors:** Nareg Pinarbasi Degirmenci, Ilknur Sur-Erdem, Vuslat Akcay, Yasemin Bolukbasi, Ugur Selek, Ihsan Solaroglu, Tugba Bagci-Onder

## Abstract

Glioblastoma is the most common type of primary brain tumor with an aggressive clinical course, and one of the cornerstones in its treatment regimen is radiotherapy. However, tumor cells surviving after radiation is an indicator of treatment failure; therefore, better understanding of the molecular mechanisms regulating radiotherapy response is of utmost importance. In this study, we generated multiple clinically relevant irradiation exposed models, where we applied fractionated radiotherapy over a long period of time and selected irradiation-survivor (IR-Surv) glioblastoma cell populations. In these cells, we examined the transcriptomic alterations, cell cycle and growth rate changes as well as responses to secondary radiotherapy and DNA damage response (DDR) modulators. Accordingly, IR-Surv cells exhibited slower growth and partly retained their ability to resist secondary irradiation. Transcriptomic analysis revealed that IR-Surv cells upregulated the expression of DDR-related genes, such as *CHK1, ATM, ATR, MGMT,* and had better DNA repair capacity as an adaptive mechanism. Separately, we report IR-Surv cells to display downregulation of hypoxic signature and the lower induction of hypoxia target genes and not exhibiting the same level of hypoxia-induced changes with naïve glioblastoma cells, as gauged by exposing cells to different hypoxia conditions. We also showed that Chk1 inhibition alone or in combination with irradiation significantly reduces cell viability in both naïve and IR-Surv cells. However, IR-Surv cells were markedly less sensitive to Chk1 inhibition under hypoxic conditions. In conclusion, consistent with previous reports, we demonstrate the utility of combining DDR inhibitors and irradiation as a successful approach for both naïve and IR-Surv glioblastoma cells as long as cells are refrained from hypoxic conditions. Thus, our findings with clinically relevant radiation survivor models will have future translational implications and benefit the optimization of combination therapies for glioblastoma patients.

## INTRODUCTION

Glioblastoma remains a significant health problem with being an incurable malignant brain tumor in adults [1]. The standard of treatment for patients diagnosed with glioblastoma has long entailed tumor resection, followed by chemotherapy and radiotherapy as described in the landmark European Organization for Research and Treatment of Cancer (EORTC) Brain Tumor and Radiotherapy Group and the National Cancer Institute of Canada [2],[3]. Recent genomic and molecular studies have shown that glioblastoma is the most heterogeneous disease among all cancer types and is composed of several cell populations with multiple genotypic origins [4]. Despite the advances in our understanding of glioblastoma genetics, cell-of-origin, or tumor heterogeneity, the survival rates have remained unchanged during the last decade.

Ionizing radiation (IR) based radiotherapy is a gold therapeutic cornerstone for glioblastoma patients. It is applied as a fractionated clinical regimen, by administering patients 2 Gy of IR over 5 days/week reaching a total of 60 Gy at the end of 6 weeks. However, despite IR and concomitantly applied chemotherapy with Temozolomide, a DNA alkylating agent, tumor recurrence is observed in majority of the patients. One of the mechanisms behind therapeutic failure is considered to be inherent or acquired therapy resistance of glioblastoma cells. The tumor cells that are radioresistant cannot be efficiently eradicated after a full dose of IR treatment suggesting that tumor cells develop adaptations to the applied therapies by undergoing genetic or epigenetic changes [5], [6]. Repopulation by IR exposed surviving glioblastoma cells during fractionated IR is among the main reasons for radiotherapy-resistant tumor recurrence [7]. To overcome this problem, several approaches for radiosensitization have been investigated, yet none of them has translated into the clinic to improve the radiosensitivity in glioblastoma patients so far [8]. Although different glioblastoma cell lines have been examined in this context, where they were exposed to short term and low doses of IR, the behavior of glioblastoma cells after long term and high dose radiation (total 60 Gy) remains largely unknown [9]-[13]. Most pre-clinical studies that interrogated the low-dose IR response of glioblastoma cell lines have shown that the radiosensitization effect is achieved by mainly targeting DNA damage repair pathways, tumor microenvironment, and cancer stem cells, but there are conflicting results with respect to obtaining radiation-persistent models in those studies [14], [15]. Currently, there are no effective therapies to target long-term IR-exposed survivor (IR-Surv) cells.

Radiotherapy induces damage to the genetic material of the cell, and affects numerous vital cellular mechanisms that may trigger radioresistance with persistent or irreparable DNA damage, activated DNA damage response (DDR), irreversible cell cycle arrest, and oncogene activation, besides several unknown reasons [16]–[18]. Furthermore, metabolic changes occur in response to IR treatment, by stimulating oxidative stress and hypoxic mechanisms. Hypoxia Inducible Factor 1 (HIF-1) stabilization or activation by IR triggers protective processes by regulating downstream target genes that can induce immunosuppressive and anti-apoptotic responses [19]. Several studies reported the improvement of radiosensitivity by blocking DDR and hypoxia pathways [20], [21]. Although targeting such pathways for glioblastoma therapy has shown promise in animal models, none has so far worked in clinical practice and improved patient survival [22], [23]. Therefore, a better understanding of IR response in clinically-relevant experimental cell models are needed to mimic radiobiological characteristics of tumors after standard clinically-applied therapeutic regimens. Moreover, growing evidence suggests the host immunity and inflammation as two conditions impacting glioblastoma progression, which clinically stratifies patients into two significant outcome groups following the same radiochemotherapy protocols, pointing out the importance of tumor stroma and microenvironment in addition to tumor characteristics [24]–[28]. As radiotherapy targets not only the tumor but also the adjacent healthy brain tissue, the inflammation and hypoxic changes in stroma and its relationship with glioblastoma require further experimental modeling to resolve related clinical discrepancy.

In this study, we established human IR-Surv glioblastoma cell models *in vitro* by exposing cells to 40-60 Gy of fractionated radiotherapy. Using established and patient derived cell lines, we selected radiation survivor cells and characterized the phenotypic and transcriptomic alterations in these cells. We demonstrated that DDR and hypoxia pathways have undergone major adaptations in IR-Surv cells in favor of improved DNA repair capacity. We showed that targeting these pathways using chemical inhibitors or switching oxygen conditions along with IR may serve as key therapeutic approaches for radiosensitization of IR-Surv cells and may be applied in the clinic to target recurrent tumors.

## RESULTS

### Generation of radiation survivor (IR-Surv) glioblastoma cell populations

To mimic the standardized radiotherapy protocol used in the clinic and generate clinically-relevant irradiation-exposed cell populations, we used three established (U373, T98G, LN229) and one primary (KUGBM8) cell line and irradiated them to a total dose of 40-60 Gy, fractionated by 2 Gy with five times a week. Parental cell lines were also passaged with irradiated samples to establish age-matched controls **(Figure 1A)**. LN229 and T98G cells could survive until a total dose of 30 Gy, and KUGBM8 cells until 40 Gy. Cell populations that survived long-term IR exposure were named as IR Survivor (IR-Surv) cells. Since U373^60 Gy^ and KUGBM8^40 Gy^ IR-Surv cells persisted longer under high exposure to IR compared to other cell lines, we mainly focused on the characterization of these and their parental pairs. The morphological analysis revealed that irradiation caused a significant increase in cell size in U373^60 Gy^ cells without affecting nucleus size **(Figure 1B, 1C)**. Compared to their parental cells, both nucleus and cell size decreased in KUGBM8^40 Gy^ cells. The percentage of multinucleated cells in the population increased in U373^60 Gy^ and KUGBM8^40 Gy^ compared to their parental pairs **(Figure 1E)**, consistent with the previous reports on HepG2 cells [29]. Long-term IR exposure also affected proliferation rates; the proliferation rate of U373^60 Gy^ cells was slower than U373 **(Figure 1F)**. However, we observed a slightly increased proliferation rate in KUGBM8^40 Gy^ cells **(Supplementary Figure S1A).** In addition, there were significant differences in the cell cycle distribution of IR-Surv cells and their parental pairs. The percent number of U373^60 Gy^ cells in the G1 phase of the cell cycle was higher than its parental pair **(Figure 1G),** but we did not observe a significant change in the cell cycle distribution of KUGBM8 cells **(Supplementary Figure S1B).**

**Figure 1.**
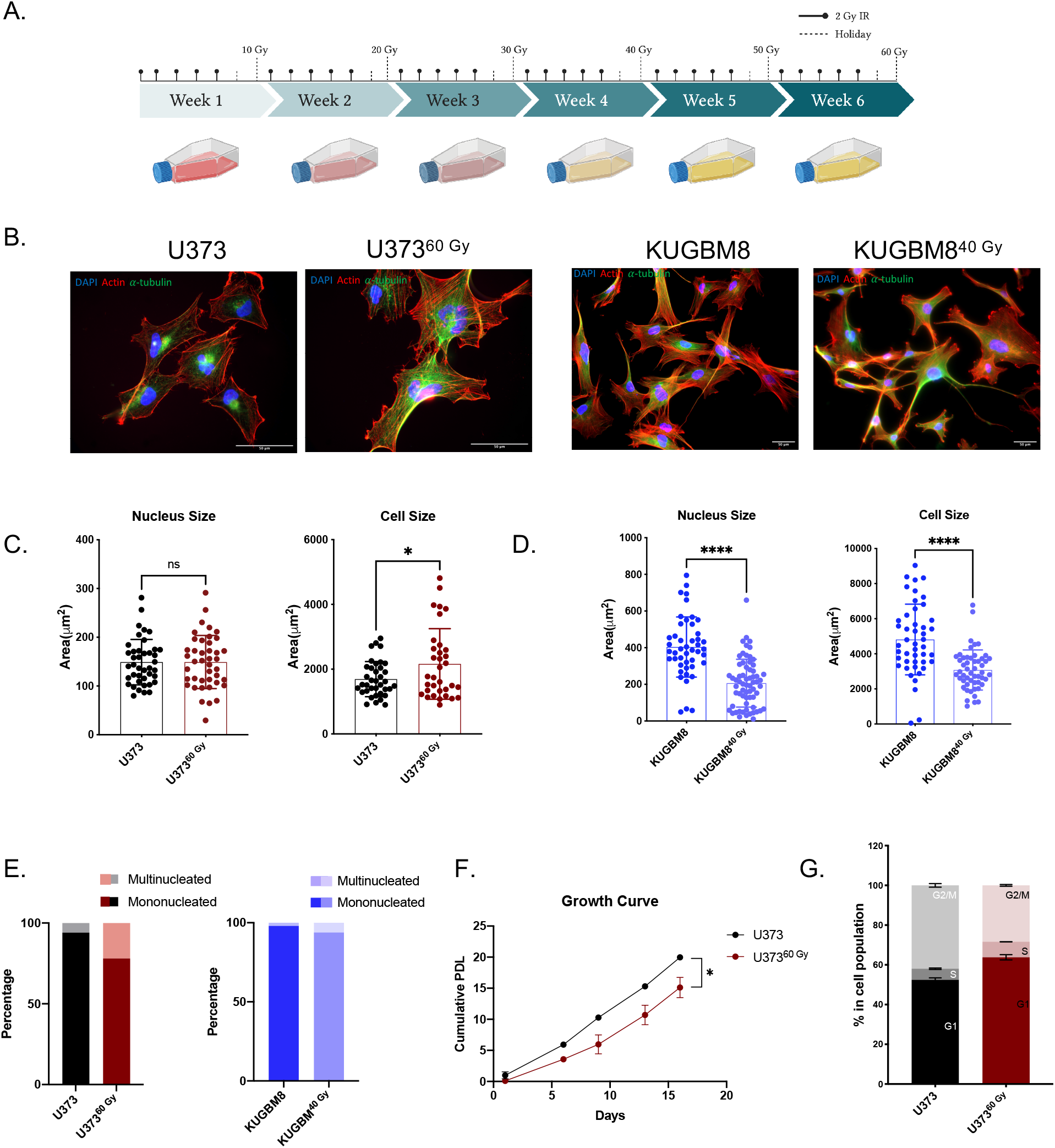
Generation of radiation survivor cell populations. **A.** Experimental setup for generation of radiation survivor (IR-Surv) cell populations (generated by Biorender.com). **B.** Immunofluorescent staining for U373 and KUGBM8 naïve and IR-Surv populations. (DAPI: blue, Actin: red, *α*-tubulin: Green, scale bar: 50 *μ*m). **C-D.** Comparison of nuclei and cell sizes between naïve and IR-Surv cells. **E.** Multinucleated cell ratios upon IR-exposure of U373 and U373^60 Gy^ cells. **F.** Proliferation rates of U373 and U373^60 Gy^ cells. **G.** Cell cycle distributions of U373 and U373^60 Gy^ cells.

To investigate whether selected IR-Surv cells maintain their persistent phenotype with secondary irradiation, we tested the viability of U373^60 Gy^ and KUGBM8^40 Gy^ cells in response to varying amounts of single-dose irradiation with clonogenic assays, which can be considered as a gold standard to assess the long-term effects of chemoradiation studies [30]. To this end, cells were exposed to 2,4,6 and 8 Gy of a single dose irradiation, and colony forming ability was measured-after 14 days **(Figure 2A)**. Accordingly, U373^60 Gy^ cells exhibited less response to radiation treatment and better colony-forming abilities than their parental pairs after 4,6 and 8 Gy treatment **(Figure 2B)**. On the other hand, we did not observe any significant difference in colony-forming abilities of KUGBM8^40 Gy^ cells **(Figure 2B, Supplementary Figure S1D)**. We also examined whether there was any cross-resistance of IR-Surv cells to Temozolomide (TMZ), the clinically applied chemotherapeutic for glioblastoma [2]. We treated naive and IR-Surv cells with increasing doses of TMZ and examine different responses of IR-Surv cells. Accordingly, U373^60 Gy^ cells had a higher IC50 value of TMZ than its parental pair; and displayed a TMz-resistant behavior (IC_50_U373 = 18.81 μM, IC_50_U373^6OG^y = 80.75 μM) **(Figure 2D).** However, KUGBM8^40 Gy^ cells displayed a better response to TMZ than their parental pair. For further elucidation of secondary therapy response, we combined single dose 4 Gy IR with high-dose TMZ. Similar to previous findings, U373^60 Gy^ cells had a higher tolerance to TMZ+IR combination, whereas KUGBM8^40 Gy^ cells were more sensitive **(Figure 2E, Supplementary Figure S1E)**. Together, we generated clinically-relevant cell line models of IR-surviving cells, one of which was derived from a well-known established cell line and the other one from a primary cell line. Despite their few differences, both IR-Surv cell lines displayed refractory behavior to secondary irradiation, mimicking the radioresistance observed in clinical settings.

**Figure 2.**
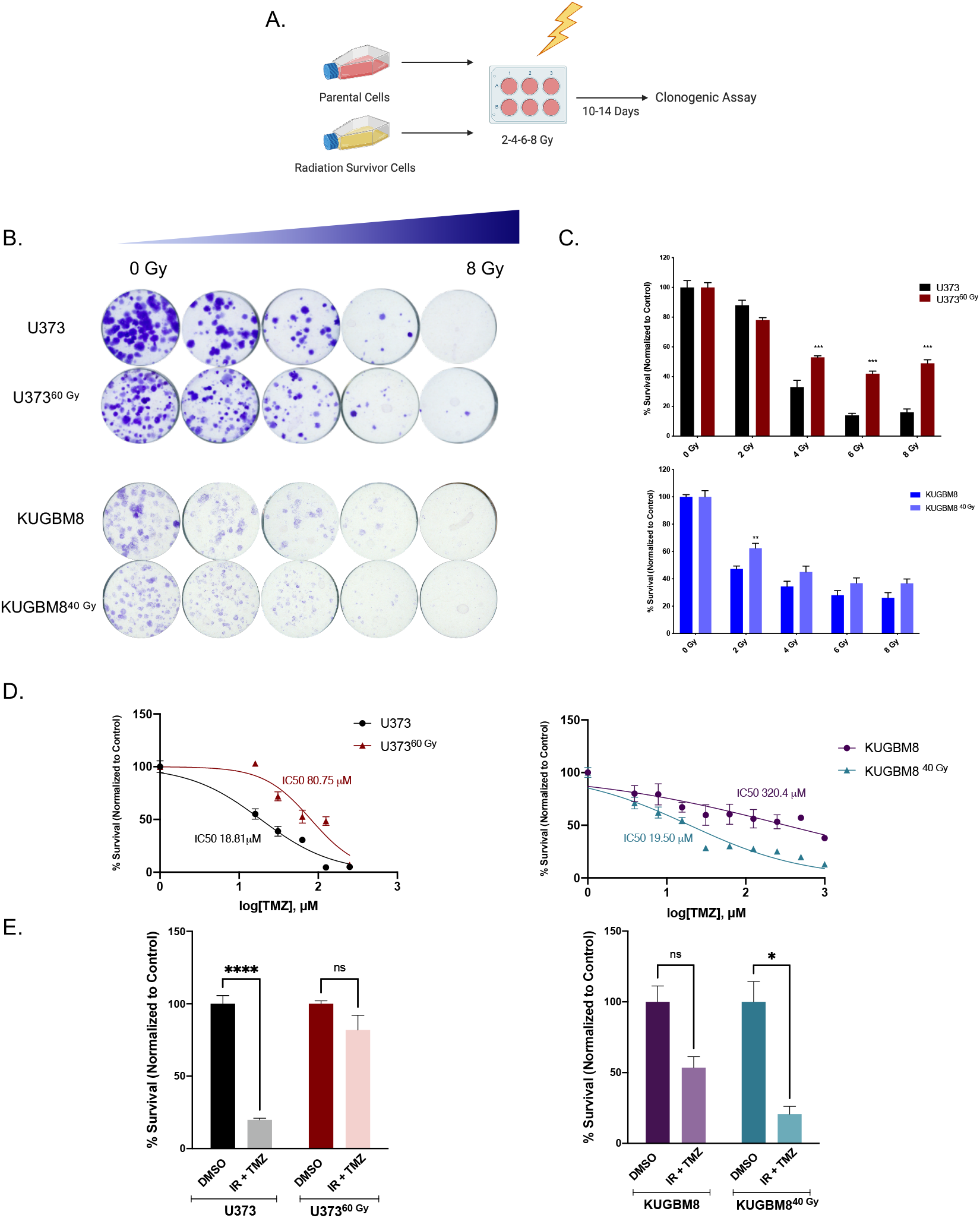
Effect of secondary IR treatment varies among different IR-Surv cells. **A.** Schematic representation of experimental set up to test the effects of secondary ionizing radiation exposure (generated by Biorender.com). **B.** Representative images of clonogenic assay of U373-U373^60 Gy^ and KUGBM8-KUGBM8^40 G^y cells upon single treatment with increasing doses of IR. **C.** Quantification of colony numbers of naïve and IR-Surv U373 and KUGBM8 cells. **D.** Dose-response curves of naïve and IR-Surv U373 and KUGBM8 cells upon TMZ treatment. **E.** Cell viabilities of cells upon TMZ and IR combination treatment.

### Transcriptomics analyses reveal changes in DNA Damage Response and Hypoxia pathways in IR-Surv cells

To understand global transcriptomic changes related to IR exposure and survival from it, RNA sequencing was performed on naive and IR-Surv U373 and KUGBM8 cell populations. Replicates from each group were clustered together in a hierarchical clustering map and Principal Component Analysis (PCA) revealed good separation of U373, KUGBM8, and their IR-Surv subpopulations from each other **(Supplementary Figure S2A, S2B).** 1346 genes were differentially expressed between U373 and U373^60 Gy^ cells; 803 of the genes were downregulated, and 543 of the genes were upregulated. These numbers were even higher between KUGBM8 and KUGBM8^40Gy^ cells; 3153 genes were downregulated, and 2141 genes were upregulated **(Figure 3A)**. To examine the differences in gene networks and pathways between parental and IR-Surv populations, we performed Gene Set Enrichment Analysis (GSEA) over 22000 identified pathways from different datasets. Pathways such as DNA Repair and double-stranded break repair were upregulated in U373 and KUGBM8 IR-Surv populations. This is not surprising, as surviving long-term exposure to ionizing radiation partly depends on adaptive mechanisms of DNA damage response and repair **(Figure 3B)** [31]. In addition, GSEA of hallmark pathways showed that pathways such as Hypoxia, ROS, and G2M Checkpoint were downregulated in IR-Surv cells **(Supplementary Figure S2C, S2D, S2E, S2F)**. Focusing on two of the activated pathways, GOBP Regulation of DNA Repair and Reactome DNA DSB Repair, we observed that the majority of the genes were upregulated in IR-Surv cells **(Figure 3C, 3D)**, suggesting IR-Surv cells rewire DNA damage recognition, response, and repair pathways to adapt to extreme IR exposure, in accordance with previous reports [32]. Interestingly, in both U373 and KUGBM8 IR-Surv populations, hypoxia-related pathways were downregulated, suggesting that cells adopt a less hypoxic signature upon IR exposure as a survival mechanism **(Figure 3B, 3E)**. To investigate the commonalities of both IR-Surv cell populations, we performed pathway analysis of shared genes between U373^60 Gy^ and KUGBM8^40 Gy^ populations on Enrichr platform using MSigDB 2021 [33]–[35]. 101 genes were commonly upregulated between IR-Surv cells; the top upregulated pathway was “IR Response Down”. 318 genes were commonly downregulated, and the top downregulated pathway was “Hypoxia” **(Figure 3F)**. These results from transcriptomic analysis provide insights into the mechanisms that IR-Surv adapt to survive IR and ultimately lead to tumor recurrence.

**Figure 3.**
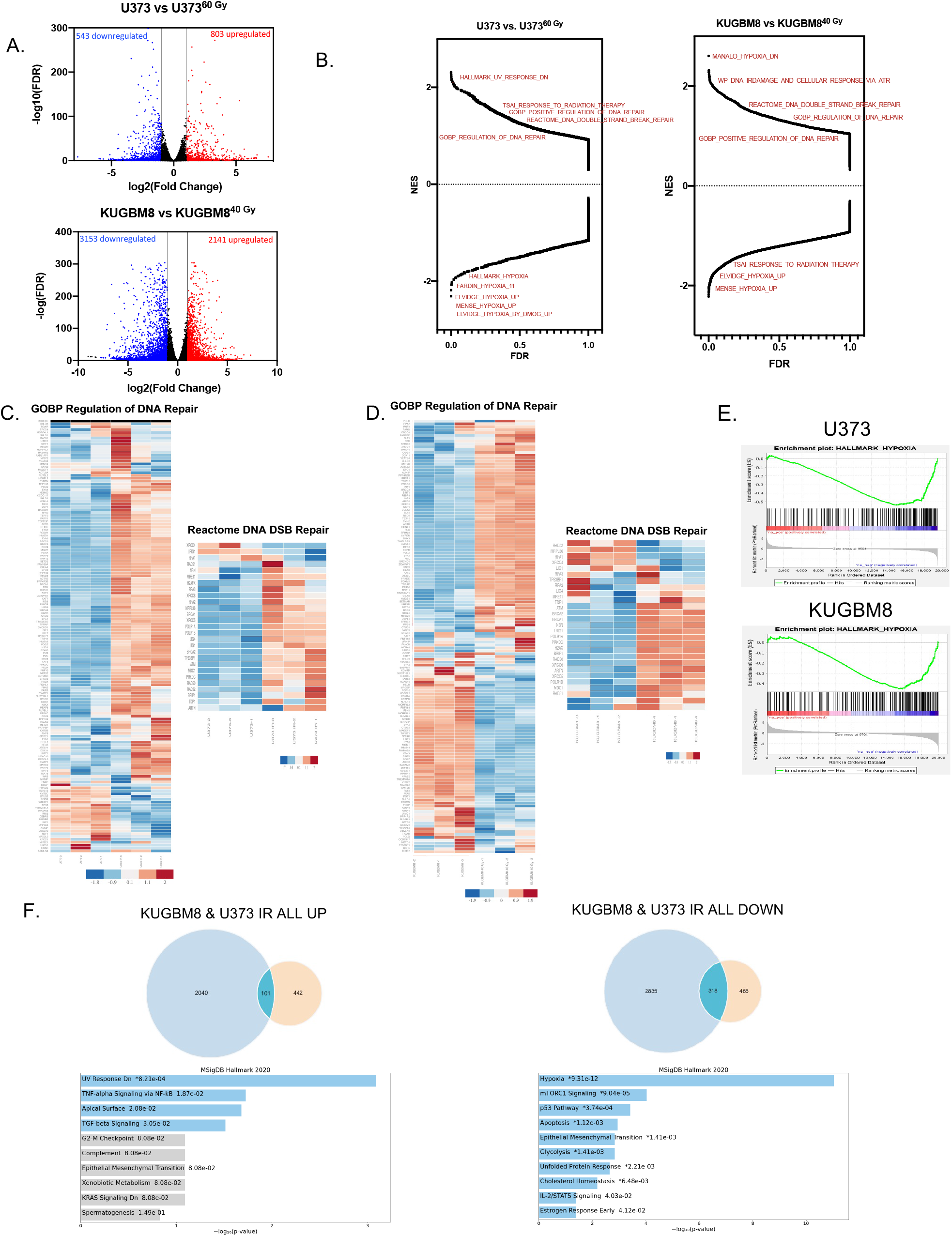
Transcriptomic alterations in IR-Surv cells. **A.** Volcano plot showing the upregulated (red) and downregulated (blue) genes in both IR-Surv U373 and KUGBM8 cells **B.** Gene Set Enrichment Analysis (GSEA) plots of U373 and KUGBM8 cells. **C-D.** Differential expression heat map of GOBP Regulation of DNA Repair and Reactome DNA DSB Repair gene sets for U373 and KUGBM8 cells. Heatmap showing z-score of log2 transformed gene expression of selected genes. **E.** Representative enrichment plots of U373 and KUGBM8 cells for Hypoxia pathway. **F.** Pathway analysis of commonly upregulated (left) and downregulated (right) genes between U373 and KUGBM8 and their IR-Surv populations.

### IR-Surv cells have higher DNA repair capacity

The overarching goal of radiotherapy is to generate DNA damage, causing genomic instability and leading to the death of tumor cells. Indeed, one major mechanism of survival from radiotherapy is through alteration of DNA damage response and repair [13]. To this end, we examined the generation and repair of DNA double-stranded breaks induced by ionizing radiation by staining for (*γ*H2AX) and Tumor suppressor p53-binding protein 1 (53BP1). After a single dose of 4 Gy irradiation of U373 and U373^60 Gy^, presence and clearance *γ*H2AX or 53BP1 levels were examined at 1 hour and 6 hours **(Figure 4A).** 4 Gy IR exposure increased 53BP1 positive-foci in both U373 and U373^60 Gy^ cells. The foci number decreased to approximately 40% in U373 cells; and to around 25% in U373^60 Gy^ at 6 hours **(Figure 4B),** indicating a different level of regulation of DBS repair by 53BP1 in IR-Surv cells. As an indicator of DNA DSB burden of cells, 1 hour after IR treatment, both U373 and U373^60 Gy^ cells had elevated *γ*H2AX-positive foci at comparable levels **(Figure 4C).** Basal levels of *γ*H2AX foci were higher in U373^60 Gy^; plausibly due to the prolonged IR exposure from which the cells survived despite DNA damage **(Supplementary Figure S3A).** After 6 hours, foci number did not change in U373 cells, but *γ*H2AX number significantly decreased in U373^60 Gy^ cells **(Figure 4C),** suggesting that IR-Surv populations had altered DNA DSB recognition and repair machinery and faster DSB break repair [36]. Furthermore, gene expression levels of several DNA damage response elements, such as *ATM, ATR, CHK1, Rad51,* and genes associated with Mismatch repair (MMR) were upregulated in U373^60 Gy^ IR-Surv cells **(Figure 4D).** This gene expression signature was not observed in KUGBM8 or other glioblastoma cell lines that were utilized to generate clinically relevant IR-Surv models **(Supplementary Figure 3B).** Expression of O6-Methylguanosine methyltransferase – *MGMT,* an important prognostic marker for glioblastoma, was upregulated both in gene and protein levels in U373^60 Gy^ cells, but not in KUGBM8^40 Gy^ cells **(Figure 2D,2E).** The protein levels of activated (phosphorylated) forms of Chk1 and Chk2, γH2AX, and Rad51 were all upregulated U373^60 Gy^ cells. However, activated or basal Atm and Atr kinase expression levels were lower in U373^60 Gy^ cells. Among MMR proteins, upregulation of Msh3 and Msh6 were observed in U373^60 Gy^ cells **(Figure 4E).** The changes in protein levels were largely consistent with RNA-seq and qRT-PCR results and highlighted an overall activated DDR state in IR-Surv cells.

**Figure 4.**
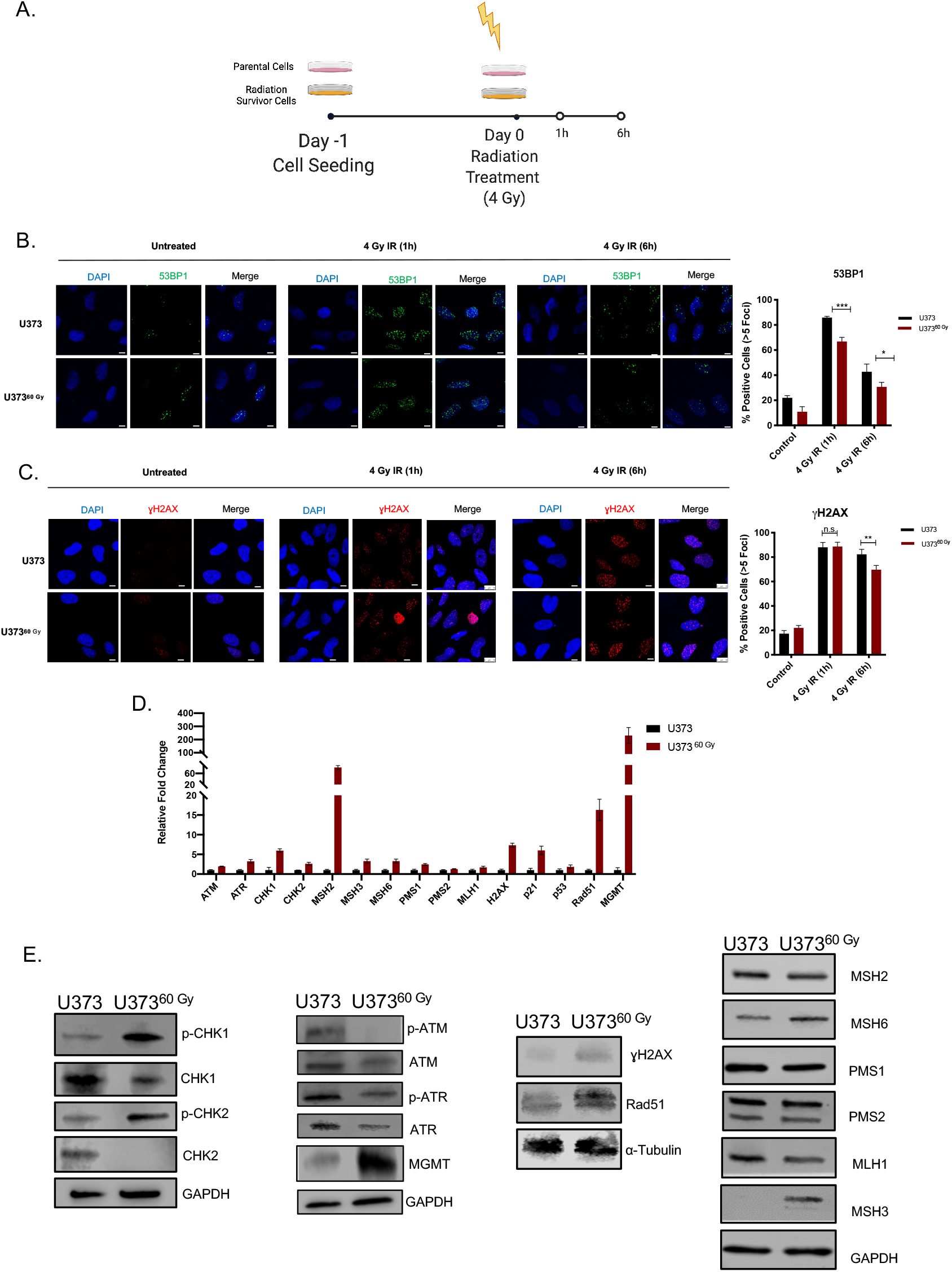
IR-Surv cells have higher DNA repair capacity. **A.** Experimental setup for immunofluorescence staining for capturing DNA damage and repair (generated by Biorender.com). **B-C.** Representative fluorescent microscope images of labelled 53BP1 and *γ*H2AX foci after 1 and 6 hours of 4 Gy IR exposure of U373 and U373^60 Gy^ (Scale bar: 10 μm) (Blue: DAPI, Green: 53BP1, Red: yH∑AX). **D.** qRT-PCR results showing expression levels of different DNA damage response and repair elements. **E.** Changes in protein levels of DNA damage response and repair elements.

### Inhibition of DDR-related kinases radiosensitizes U373 and U373^60 Gy^ cells

DNA damage response and repair pathways are among the most targeted pathways for therapeutic purposes in cancer. Inhibition of central regulators of DDR, such as Atm, Atr, Chk1, and Chk2 is considered a prime therapeutic approach in chemo- or radiosensitization studies [37]-[39]. Based on our observations with IR-Surv cells, which activated DDR to adapt to long-term IR, we interrogated whether their inhibition would sensitize IR-Surv cells to irradiation. We selected five DDR-related kinase inhibitors (DDRi) (AZD7762, AZD6738, KU55933, BML-277, and LY2603618) targeting Atm, Atr, Chk1, or Chk2 **(Figure 5A)**. U373 and U373^60 Gy^ both responded to DDRi in a dosedependent manner **(Figure 5A, 5B).** Further, DDRi radiosensitized both U373 and U373^60 Gy^ cells, but the degrees of sensitization were different between them, when examined with short-term viability assays **(Figure 5C)**. In addition, we tested the effect of DDRi and IR combination treatment on a long-term (14-day long) clonogenic assay. Accordingly, U373^60 Gy^ cells were slightly less responsive to KU55933 (Atm inhibitor) individual treatment, consistent with our initial findings. Combination treatment of DDRi and single-dose 4 Gy IR were very effective on both U373 and U373^60 Gy^ cells **(Figure 3D,3E).** These results suggest IR-Surv cells with increased DNA damage response activity, can be sensitized to IR treatment using DDR inhibitors.

**Figure 5.**
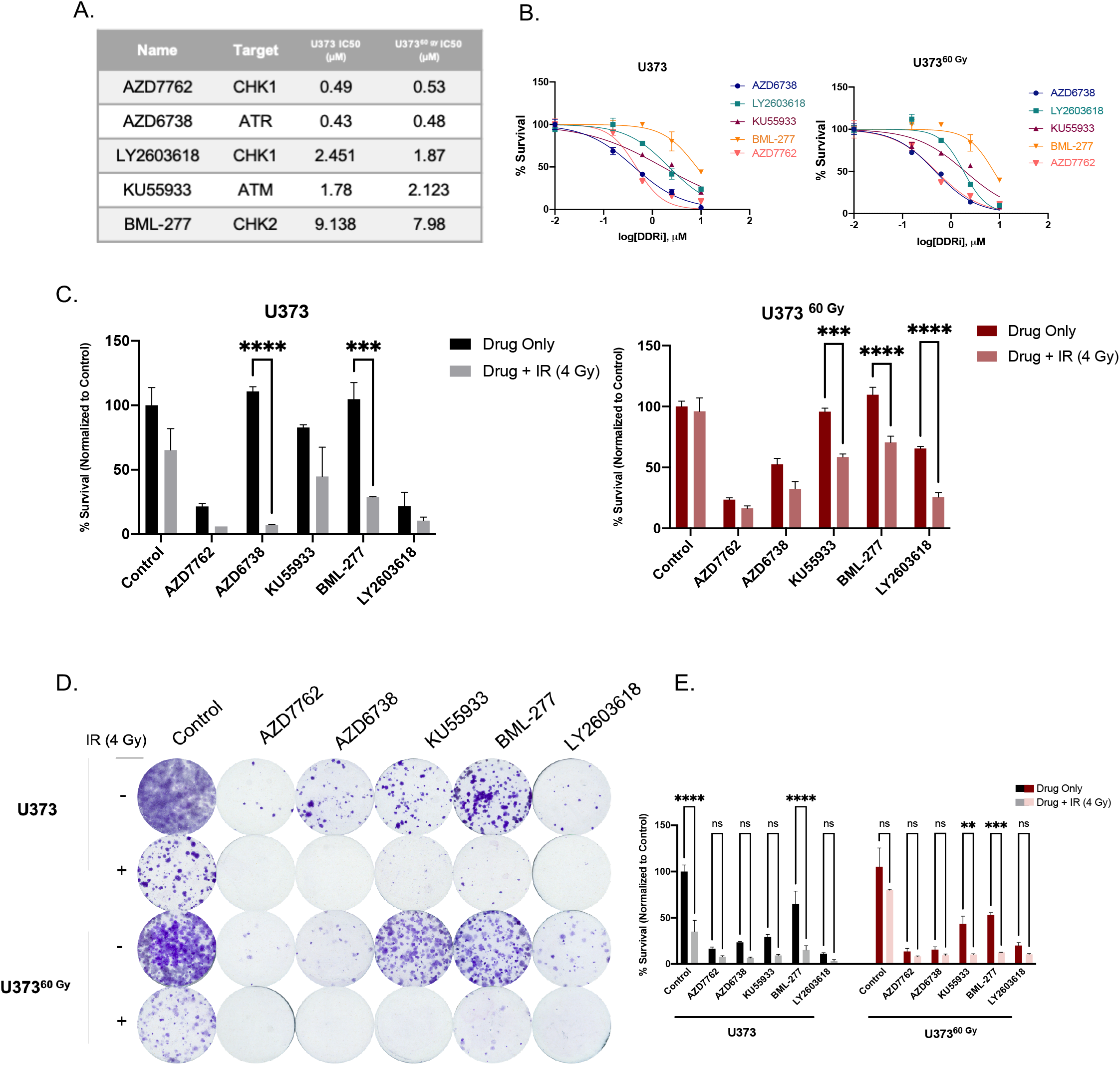
Inhibition of DDR-related kinases radiosensitizes U373 and U373^60 Gy^ cells. **A.** List of inhibitors used and IC50 values of U373 and U373^60 Gy^ cells. **B.** Doseresponse curves of inhibitors of DDR-related kinases. **C.** Cell viability results of short-term treatments of DDRi + IR combination on U373 and U373^60 Gy^ cells- **D.** Representative images of clonogenic assay of combination treatment of DDRi and single dose 4 Gy radiation on U373 and U373^60 Gy^ cells. **E.** Quantification of colony numbers of DDRi + IR combination treatments.

### IR-Surv cells have lower hypoxic state and exhibit poor response to DDR inhibition under further hypoxia

As hypoxia was identified as the top downregulated pathway from our transcriptomic analysis, we investigated the behavior of IR-Surv cells by exposing them to hypoxic conditions. For this, we examined three conditions, control (normoxia), acute hypoxia (applied for 1 day) and chronic hypoxia (applied for 4 days) **(Figure 6A)**. As a mimic for irradiation, we used DSB causing drug, Bleomycin [40], and investigated the activation of H2AX under normoxic and hypoxic conditions upon 2 hours of Bleomycin treatment. Accordingly, under normoxic conditions, Bleomycin increased γH2AX activation in both U373 and U373^60 Gy^ cells significantly. The γH2AX activation levels were similar under hypoxia in U373 cells. However, U373^60 Gy^ cells exhibited slightly less γH2AX activation under hypoxic conditions, suggesting a different mode of adaptation to DNA damage and repair **(Figure 6B)**. These adaptations have possibly affected cell cycle progression. Chronic hypoxia did not alter cell cycle distribution of U373^60 Gy^, only affecting U373 cells through G2/M arrest (**Figure 6C).** Downregulated response to hypoxia was also observed at gene expression level. Upon hypoxia, IR-Surv cells did not upregulate vascular endothelial growth factor *(VEGF),* which can be considered a biomarker for hypoxia [41], as much as parental cells did. In addition, *CHK1* upregulation in U373^60 Gy^ cells was not observed to the same extent as in parental U373 cells **(Figure 6D)**. To test whether the cells’ response to DDRi would be altered under hypoxic conditions, we treated U373 and U373^60 Gy^ with low-dose Chk1 inhibitors, AZD7762 and LY2603618. To highlight their effects on cell viability, we applied low doses of both AZD7762 and LY603618 under normoxic and hypoxic conditions to both U373 and U373^60 Gy^ cells. While 4 days of Chk1i treatment was very effective on U373 cells, it became more effective under hypoxic conditions. However, similar cell viability was observed for U373^60 Gy^ cells under both normoxic and hypoxic conditions **(Figure 6E).** To test whether this phenotype is temporary or exclusive to short-term hypoxia, we performed a 14-day clonogenic assay in hypoxic conditions **(Figure 6F).** After Chk1i treatment, cells were incubated in a hypoxic incubator for 14 days. Clonogenic assay results revealed that U373^60 Gy^ cells were far less sensitive to Chki-hypoxia combination treatment than U373 cells **(Figure 6G)**. Thus, IR-Surv cells exhibit resistance to Chk1 inhibition, the sensitization can be achieved through IR combination, but not to sufficient degrees in hypoxic conditions.

**Figure 6.**
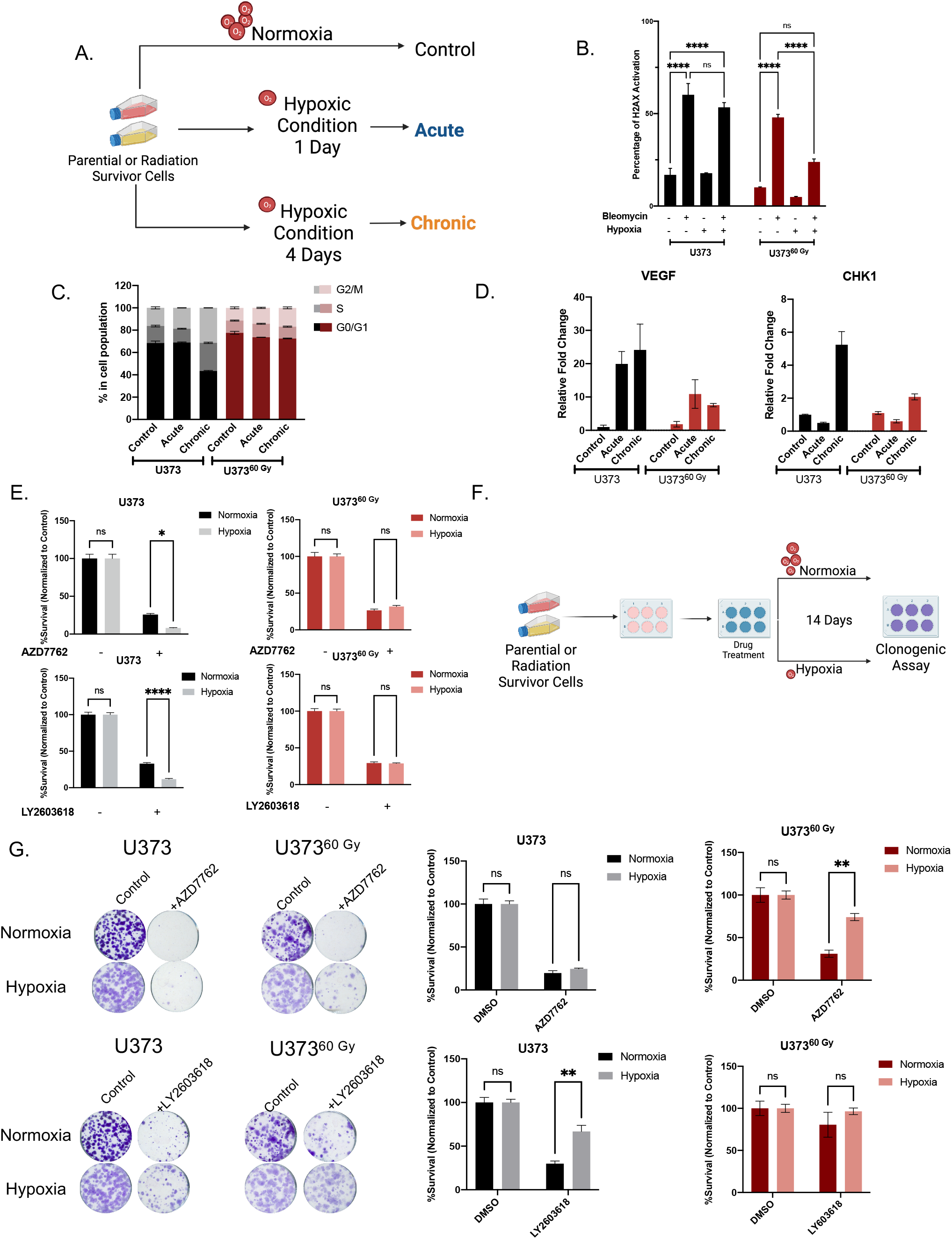
IR-Surv cells have poor DDRi response in hypoxic conditions. **A.** Schematic representation of experimental flow in hypoxic conditions (generated by Biorender.com). **B.** Assessment of H2AX activation in combination treatment of hypoxia and Bleomycin. **C.** Cell Cycle distribution of naïve and IR-Surv cells in normoxia and hypoxia. **D.** Gene expression levels of *CHK1* and *VEGF* upon culturing in acute and chronic hypoxia. **E.** Cell viability differences upon treatment of CHK1 inhibitor AZD7762 and LY2603618 in normoxia and hypoxia. **F.** Schematic representation of experimental setup for colony formation assay in long-term hypoxia. **G.** Representative clonogenic assay images and quantification of U373 and U373^60 Gy^ cells upon treatment of CHK1 inhibitor AZD7762 and LY2603618 in normoxia and hypoxia.

## DISCUSSION

As high proliferation and infiltration capacity, the ability to adapt and develop resistance to therapies are significant hallmarks of the glioblastomas [42], the main treatment options of surgery, chemotherapy, and radiotherapy seem not enough for cure. Despite the refined RT regimens and TMZ administration, recurrence occurs mostly in central high dose radiotherapy field within 90% of patients due to intrinsic or acquired therapy resistance of tumor cells [5], [43]–[45]. Therefore, understanding the molecular mechanisms behind this adaptive persistence is of utmost priority to design effective therapeutic strategies. In this study, we employed a clinically relevant radiotherapy regimen to investigate the phenotypic alterations of surviving glioblastoma cells to recapitulate the early stages of recurrence, and demonstrated that IR-Surv cells had increased DNA damage repair capacity and reduced response to hypoxia, while documented for the first time in literature that downregulation of hypoxic signature as well as the lower induction of hypoxia target genes, through functional assays and transcriptomic analysis.

By utilizing DNA damage response-related kinases, we showed that IR-Surv cells have a slight resistance to DNA damage response-related kinase inhibition, but these cells can be sufficiently eradicated by using inhibitors combined with single-dose IR exposure. Furthermore, we showed that IR-Surv cells reclaim resistance to Chk1 inhibition in hypoxic conditions. Together, our results suggest that IR-Surv cells may become more resistant to DDR inhibition in long-term hypoxic conditions **(Figure 7)**, which provides insight into the future design of effective combinatorial radiotherapy strategies.

**Figure 7.**
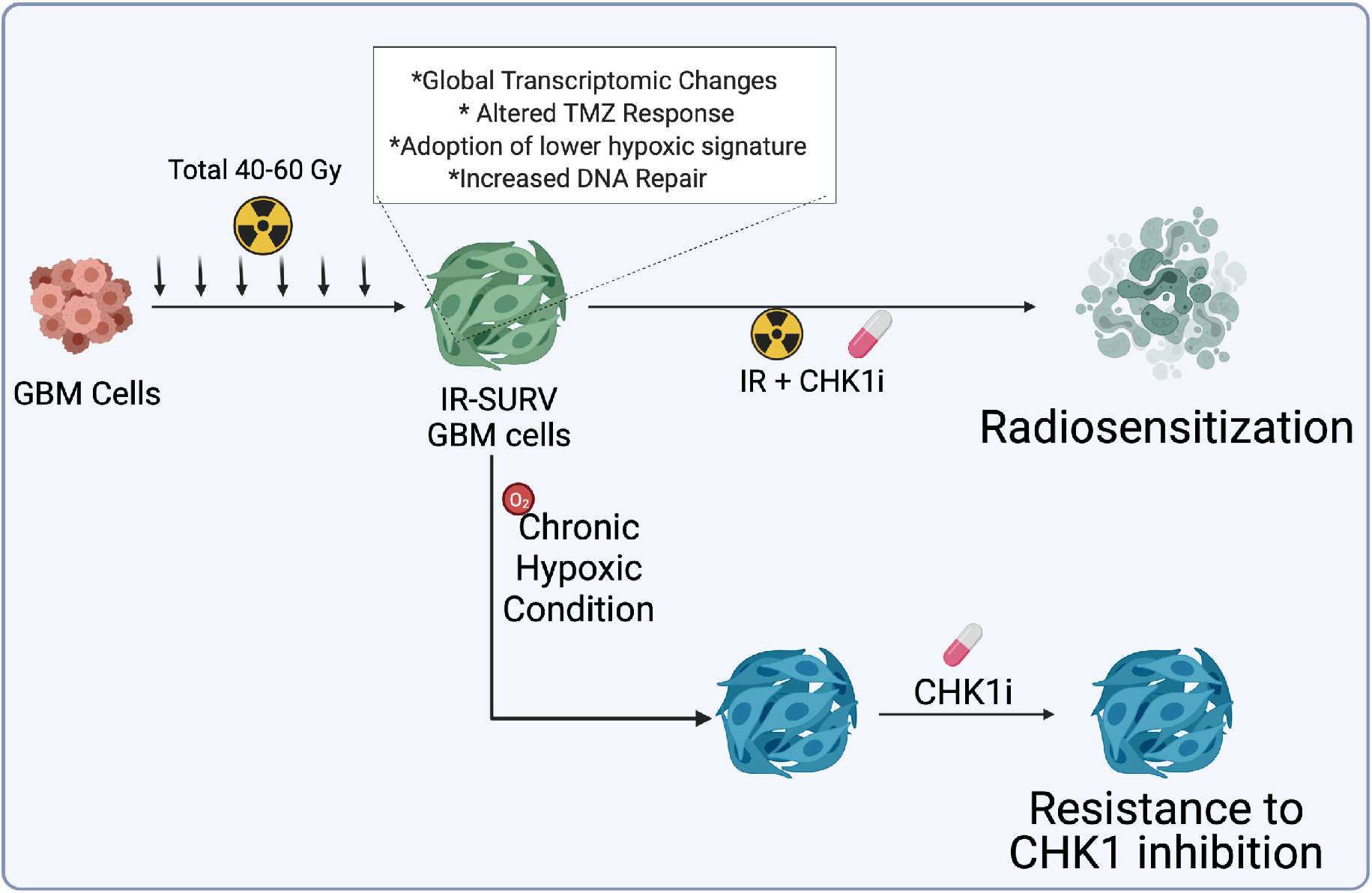
Graphical abstract of the study.

The radiation response and mechanisms of radioresistance are extensively studied in different cancer types, both primary and recurrent models. The model we exploit mimics the radiotherapy schedules applied to patients, which was previously used in few studies [46], [47]. In the generation of radiation-survivor cell lines, we tried to remain faithful to the “Clinically Relevant Radioresistant (CRR) cell line” notion [48]. CRR concept in the generation of radioresistant cancer model is based on conventional fractional RT, exposure to 2 Gy IR once a day for more than a month. The cells were exposed to 2 Gy IR/day for five days; the fraction between two treatments was two days. In those fractioned days, cells were passaged along with parental cells to ensure cells were not under stress besides radiation therapy. This way, we succeeded in generating age-matched parental/radiation-survivor (IR-Surv) cell line pairs. To examine the survival capacities of glioblastoma cell lines with different genetic landscapes, we utilized 1 primary cell line generated in our laboratory, and 3 established cell lines [49]. However, the endurance levels of cell lines to total IR treatment were variable. The highest dose exposure was achieved by U373 cells (60 Gy), followed by KUGBM8 (40 Gy), and T98G and LN229 cells (30 Gy). The diversity of endurance of the cells may depend on various factors yet to be discovered. From the analysis of common mutations in glioblastoma, PTEN and p53 status could be among indicators of radiation persistence [50]. In the case of PTEN, where KUGBM8 and LN229 are wild-type, T98G cells have missense mutation. On the other hand, U373 have null mutation of PTEN, correlating our observed radiation-survivor phenotype with different studies suggesting PTEN loss or mutation leading to radioresistance [51]. Except for KUGBM8 cells, all the cells used in the study were p53 mutant. Like most glioblastoma tumors and established cell lines, radiation response depends not solely on one gene or a pathway but on various components of tumor progression [49], [51].

As a representative model, we chose one established cell line and one primary cell line to investigate radiation-induced alterations. Although U373^60 Gy^ and KUGBM8^40 Gy^ share common characteristics such as increased ratio of multinucleated cells and retained colony formation abilities after secondary IR exposure, they exhibit different phenotypes and therapy responses. We have not observed cell cycle arrest at the G1 phase in KUGBM8^40 Gy^ cells, which is reported as an indicator for radioresistance [38]. Cells may have undergone senescence after radiation as defense mechanism, in consistency with different reports [52]. In addition, response to TMZ and adjuvant TMZ+IR also differed between U373^60 Gy^ and KUGBM8^40 Gy^ cells, very likely due to differential MGMT expression levels of these IR-Surv cells. In consistent with high MGMT expression being related to poor response to TMZ [53], IR-Surv cells that upregulated MGMT displayed collateral resistance to TMZ. This suggests that some tumors may respond to TMZ when they are in a naïve state prior to IR treatment, however prolonged IR treatment may cause the survivor tumor cells to exhibit cross-resistance to TMZ, making the chance of recurrence higher.

Our RNA sequencing analyses highlight differentially expressed pathways in IR-Surv cells. Despite U373^60 Gy^ and KUGBM8^40 Gy^ displaying few differences in functional assays, they share common features in the context of altered pathways upon long-term IR exposure. As the overarching goal of radiotherapy is to generate DNA damage directly or indirectly [31], [54], our results stand in parallel that IR-Surv cells show notable alterations in DNA damage response and repair pathways. With increased genomic instability, IR-Surv cells alter DNA repair machinery to survive and adapt to constant exposure to IR via increased expression of DNA damage response and repair-related proteins. This adaptation partly relies on increased DNA repair capacity. Radiation-exposed cells also survive with a higher burden of genomic instability, which is not elucidated in our study but prompts future work [55], [56]. Our observations suggest that U373^60 Gy^ cells have a higher burden of DNA damage, yet upon further IR-exposure, they have higher DNA damage clearance and repair capacity than their naive pair. We utilized a panel of inhibitors targeting DNA damage response-related kinases for radiosensitization. These DDRi inhibitors were studied in different cancer types in various conditions [39], [57], [58]. Our study showed that individual DDRi treatments were not sufficient for eradication of IR-Surv cells, suggesting that increased DNA repair capacity compensates for inhibition of these kinases.

Ataxia-Telangiectasia Mutated (Atm) kinase is activated explicitly upon DSB generation, which activates downstream effector kinases Checkpoint Kinase 1 (Chk1) and Checkpoint Kinase 2 (Chk2) [59]. The inhibitors of Atm, Chk1, and Chk2 kinases are extensively studied, combined with chemotherapeutic drugs and radiotherapy [20], [60], even though multiple clinical trials are currently proceeding. As involved in regulating DDR and cell cycle checkpoints, Chk1 is one of the ideal targets for radiosensitization [61], [62] Our study correlates with radiosensitization studies in the context of Chk1 function, as radiation-mediated Chk1 activation can be exploited for radiosensitization of IR-Surv cells. On the other hand, a phase I trial of AZD7762 with irinotecan in glioblastoma was stopped because of toxicity reports (NCT00473616). Therefore, there is a need for the development of new therapeutic strategies with new-generation DDR inhibitors and radiotherapy with lower cytotoxicity and improved efficacy. Our study exemplifies a treatment model combining a low dose of DDRì and radiotherapy in both naive and IR-Surv cell populations.

Therapy resistance of the tumors does not solely depend on the genetic and epigenetic landscape of tumor cells but is highly influenced by the tumor microenvironment [63]. Hypoxia is one of the highly studied concepts in the field of radiobiology, associated with all 6 R’s (Repair, Redistribution, Repopulation, Reoxygenation, Reactivation of immune response, and Radiosensitivity) [64]. Indeed, the tumor core, which is highly hypoxic, is known to be more radioresistant [65], [66]. Decreased oxygenation of tumor cells makes radiotherapy ineffective as ionizing radiation fails to generate reactive oxygen species that would lead to DNA damage [67]. On the other hand, the opposite scenario is still not elucidated. In this study, we showed that IR-Surv cells exhibit downregulation of hypoxic signature through transcriptomic analysis and by the lower induction of hypoxia target genes, such as *VEGF.* We showed that cells that escape from chronic radiation did so by several adaptive changes, one of which resulted in a lower hypoxia response. This might seem in contrast with the knowledge in radiobiology at first sight. However, this finding, which is based on two different IR-Surv models, may suggest that tumor cells can find a way to counteract the high pressure exerted by chronic IR and become more vulnerable at the end. IR-Surv cells’ response to DDR inhibition, particularly to Chk1 inhibition was also markedly less under hypoxia in IR-Surv cells [68], suggesting that further applications of DDRi need to take into account the hypoxic nature of tumors for best clinical translation.

Taken together, we generated useful clinically relevant radiation survivor models that exhibit several major adaptive mechanisms. We showed that efficacy of radiotherapy not only depends on hypoxic conditions, but also irradiation-escaped cells may display resistance to hypoxia. In addition, targeting DDR kinases such as Chk1 is effective on irradiation-escaped cells but the efficacy would decrease under hypoxia. These results could provide insight into designing effective treatment strategies for recurred tumors from radiotherapy.

## Materials and Methods

### Cell Culture and Reagents

Glioblastoma cell lines U373, LN229, and T98G were available from the American Tissue Type Culture Collection (USA). KUGBM8 primary cell line was established by Dr. Filiz Şenbabaoğlu from patient samples in collaboration with Koç University Hospital Neurosurgery Department [49]. Protocol of primary cell line generation was adapted from [69]. All parental and irradiated cell populations cells were cultured in DMEM (Gibco, USA) supplemented with 10% FBS (Gibco, USA) and 1% Penicillin-Streptomycin (Gibco, USA). All cells were maintained at 37°C in a humidified incubator with 5% CO2. To achieve hypoxic conditions, cells were maintained at 37°C in a humidified incubator with 5% CO2 and 1% O2. AZD7762 (Selleckchem, S1532), AZD6738 (Ceralasertib, Selleckchem, S7693) LY2603618 (Rabusertib, Selleckchem, S2626), KU55933 (Selleckchem, S1092), BML-277 (Selleckchem, S8632), Bleomycin (Selleckchem, S1214) and Temozolomide (Selleckchem, S1237) were used for drug treatment experiments.

### Generation of Radiation Exposed Cell Lines

Cells were irradiated with 6MV X-Ray at a dose rate of 600MU/min in Varan iX model linear accelerator, located at the Radiation Oncology Department of Koç University Hospital. To mimic clinically relevant standardized radiotherapy, cells were exposed to 2 Gy IR every day for 4-6 weeks. U373 cell lines were exposed to 60 Gy ionizing radiation (IR) for 6 weeks. Irradiation of T98G, LN229 cell lines were concluded at 30 Gy, and KUGBM8 cell irradiation was completed after 4 weeks (40 Gy).

### Clonogenic Assay

All parental and irradiated cells were seeded as 750 cells/well to 6-well plates as triplicates and exposed to single doses of ionizing radiation of 2-4-6-8 Gy for each plate and incubated for 14 days. Wells were washed with 1x DPBS twice, and colonies were fixed with ice-cold methanol treatment for 5 minutes. After fixation, wells were washed with 1x DPBS twice and incubated with crystal violet for 15 minutes. Crystal violet was removed, and plates were washed. After the plates were dried, plates were scanned, colony densities for each well were quantified with Adobe Photoshop CC 2019 (USA).

### Cell viability assay

Cells were seeded as 1000 cells/well, treated with the corresponding drug on day 1, and/or exposed to 4 Gy IR treatment on day 3. On day 5, MTT solution (3 mg/ml) was added as 25 μl per well and incubated for 4 hours at 37°C. After incubation, culture medium was aspirated, 100 μl DMSO was added and dissolved. Plate reading was performed in Synergy H1 Reader (Biotek, USA) at 570 nm wavelength.

Survival was described as a percentage of viable cells of each sample compared with DMSO control groups.

### Cell Cycle Assay

Cells were harvested from 6-well plates, and pellets were washed with ice-cold PBS. For fixation, pellets were resuspended with cold ethanol (70%) and incubated for 30 minutes at 4°C. After fixation, samples were centrifuged and washed with PBS twice, resuspended in 50 μl PBS containing RNase A (100 μl /ml) and incubated at room temperature for 15 minutes. Propidium Iodide (PI) (50 μl/ml) was added, and samples were incubated at room temperature for 30 minutes. Stained samples were analyzed by BD Accuri C6 (BD Biosciences, USA) flow cytometer, and 10.000 events were recorded for each sample. For cell cycle analysis in hypoxic cells, The Muse^®^ Cell Cycle Kit (MCH100106) was used according to the manufacturer’s protocol.

### H2AX Activation Assay

Cells were harvested from 6-well plates following the corresponding treatment, and H2AX activation was quantified with The Muse^®^ H2A.X Activation Dual Detection Kit (MCH200101) according to the manufacturer’s protocol.

### Western Blotting

Cell pellets were lysed in an appropriate volume of lysis buffer (1% NP40, 150 mM NaCl, 1mM EDTA, 50 mM Tris-HCl (pH 7.8), 1 mM NaF) containing 0.1 mM PMSF, and 1X protease inhibitor cocktail (complete Protease Inhibitor Cocktail Tablets, Roche). Following 30 minutes of incubation in the lysis buffer, the lysates were sonicated and centrifuged (12000 rpm, 4°C, 15 min). Samples were denatured in 4xSDS sample buffer at 95°C for 5 minutes. For equal protein loading, Pierce™ BCA (Bicinchoninic Acid) Protein Assay (Thermo Fisher) was performed, and calculations were done accordingly. For immunoblotting, equal amounts of protein were separated by SDS-polyacrylamide gel electrophoresis and transferred onto a PVDF membrane by Trans-Blot^®^ TurboTM RTA Mini PVDF Transfer Kit (Biorad, USA). Later, the membranes were blocked with 5% non-fat dry milk in TBS-T (20 mM Tris-HCl, pH 7.8, 150 mM NaCl, 0.1%, v/v Tween-20) at RT for 1 hour. After blocking, the membrane was incubated with primary antibodies overnight (4°C). The list of primary antibodies is listed in **Supplementary Table S2.** The membrane was washed three times with TBS-T for 15 min. The corresponding appropriate horseradish peroxidase coupled secondary antibodies (Cell Signaling, 1:10.000) were incubated for 1 h, and the membrane was washed three times with TBS-T. Blots were incubated with ClarityTM Western ECL Substrate (Biorad, USA) and visualized using an Odyssey Scanner (LiCor Biosciences, USA).

### Quantitative Real-Time PCR (qRT-PCR)

To determine respective mRNA expression, parental and irradiated cell pellets were collected, and RNA isolation was performed with NucleoSpin. RNA Isolation Kit according to manufacturer’s instructions (Macherey-Nagel). RNA concentrations were measured with Nanodrop. With reverse transcriptase reaction, 900 ng of cDNA was obtained using M-MLV Reverse Transcriptase (Invitrogen). mRNA expression levels of specific genes were detected by LightCycler. 480 SYBR Green I Master (Roche). Sequences of used primers are listed in **Supplementary Table S1**.

### Immunofluorescence Staining

Cells were seeded on 24-well plates on coverslips as 20.000 cells/well. After irradiation, the media was removed, and cells were washed with PBS twice. Cells were fixed with 4% PFA for 5 minutes at room temperature. After PBS wash, fixed cells were treated with 0.1% Triton X-100 for 5 minutes and washed with PBS. Each well was treated with 250 μl SuperBlock IHC Blocking Solution (ScyTek Laboratories, USA) at room temperature for 15 minutes. After rinsing wells with PBS, coverslips were incubated with primary antibodies overnight at 4°C. Coverslips were washed with PBS again and incubated with respective secondary antibodies at room temperature for 1 hour in the dark. The list of primary and secondary antibodies is given in **Supplementary Table S3**. Images were taken at Zeiss Axio Imager M1 (Germany) at 40x magnification. Foci numbers were counted for each condition and normalized to untreated control groups.

### RNA-Sequencing and Analysis

Total RNAs of irradiated and parental cells were isolated using Macherey-Nagel NucleoSpin^®^ RNA Isolation Kit. Based on protocols of BGISEQ-500 platform, RNA-seq libraries were prepared. Libraries were sequenced on a BGI seq 500 platform using 20 million single-end reads per sample. For U373 and U373^60 Gy^ and KUGBM8 and KUGBM8^40 Gy^, independent 3 replicate samples were sent for sequencing. Sequenced data were converted to FASTQ files using BGISEQ-500 platform at BGI Genome Sequencing Company (Beijing, China). FASTQ files were uploaded into Genialis Expressions Platform (Texas, USA) and analyzed. According to differential expression data, volcano plots were generated, with log2(FC)=1 and 0.05% FDR cut-offs. Also, heat map representations were used for selected gene sets to visualize differential expressions. Pathway analysis was performed with Enrichr gene list enrichment analysis tool directly linked to Genialis website. Gene Set Enrichment Analysis was performed. The RNA-seq data have been deposited in NCBI’s Gene Expression Omnibus (GEO), with accession number GSE199862.

### Statistical Analysis

All normalizations were performed on non-radiated or untreated samples, denoted as 100% using GraphPad Prism version 9.0 (USA) and Microsoft Excel 2018. Significance analysis was performed with student’s t-test and two-way ANOVA (n.s denote not significant, for p-values, *, ** and *** denote p < 0.05, p < 0.01 and p < 0.001 respectively, two-tailed Student’s t-test).

## Author Contributions

Study design: T.B.O., I.S.E., and N.P.D.; methodology: N.P.D, T.B.O, U.S, Y.B., I.S.; data generation and analysis: N.P.D., I.S.E., V.A.; data interpretation: T.B.O., N.P.D, V.A, I.S.E., U.S., Y.B. drafted the manuscript: T.B.O., N.P.D., I.S.E; approved final manuscript: all authors.

## Acknowledgments

We thank Yakup Barkodat, Esra Serbest Erkan, Mert Topçu, and Ali ihsan Atasoy at Koç University Hospital, Department of Radiation Oncology for assisting with irradiation protocols.

