## Supplemental Figures for "Chronically radiation-exposed survivor glioblastoma cells display poor response to Chk1 inhibition"

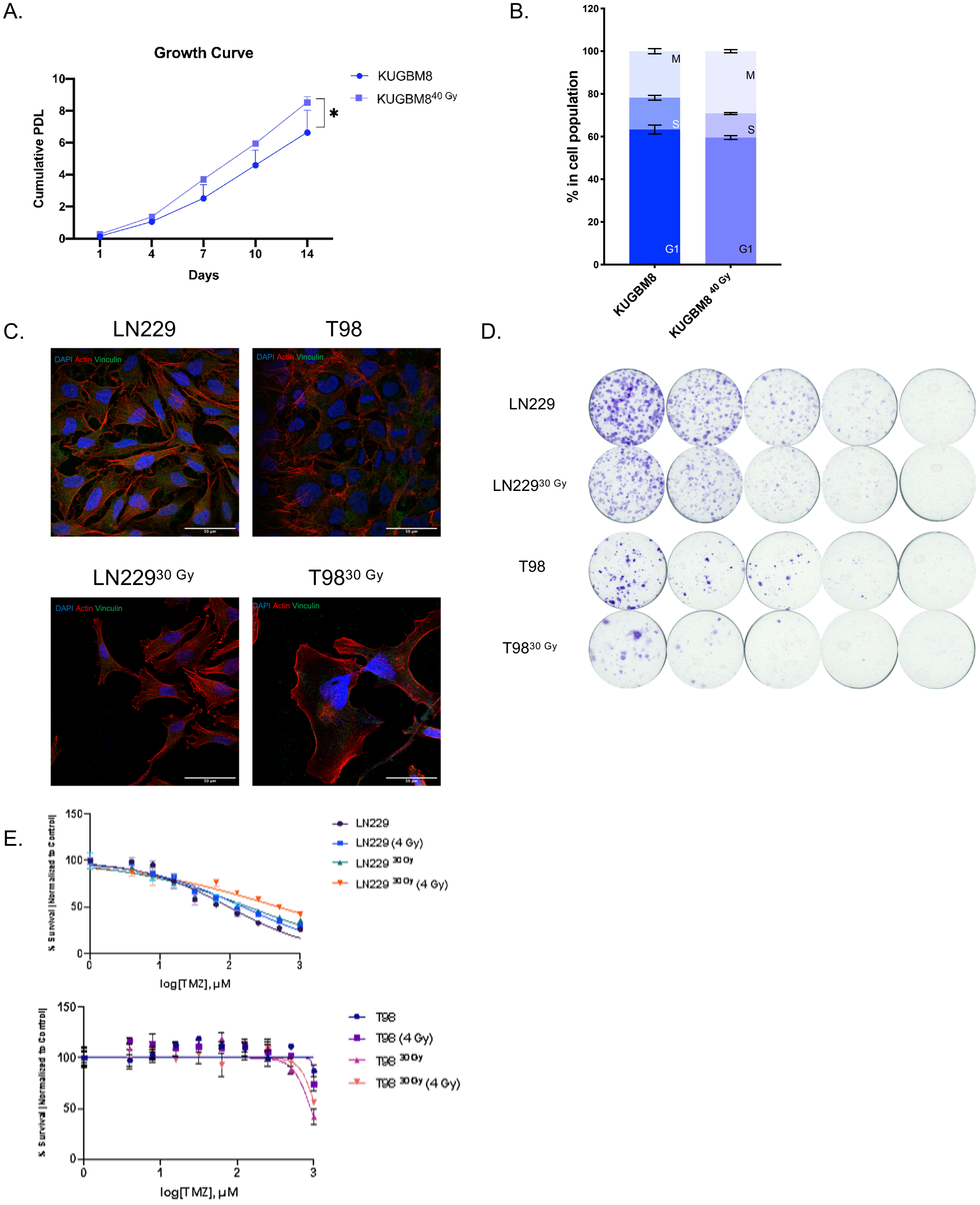


**Supplementary Figure S1.** A. Proliferation rates of KUGBM8 and KUGBM8$^{40\ Gy}$ cells. B. Cell cycle distributions of KUGBM8 and KUGBM8$^{40\ Gy}$ cells. C. Immunofluorescent staining for T98 and LN229 cells exposed to IR (DAPI: blue, Actin: red, Vinculin: Green).. D. Representative images of clonogenic assay upon increasing dose of IR of T98 and LN229 cell lines. E. Dose-response curves of naïve and IR exposed T98 and LN229 cells upon TMZ and IR only treatment, and in combination.

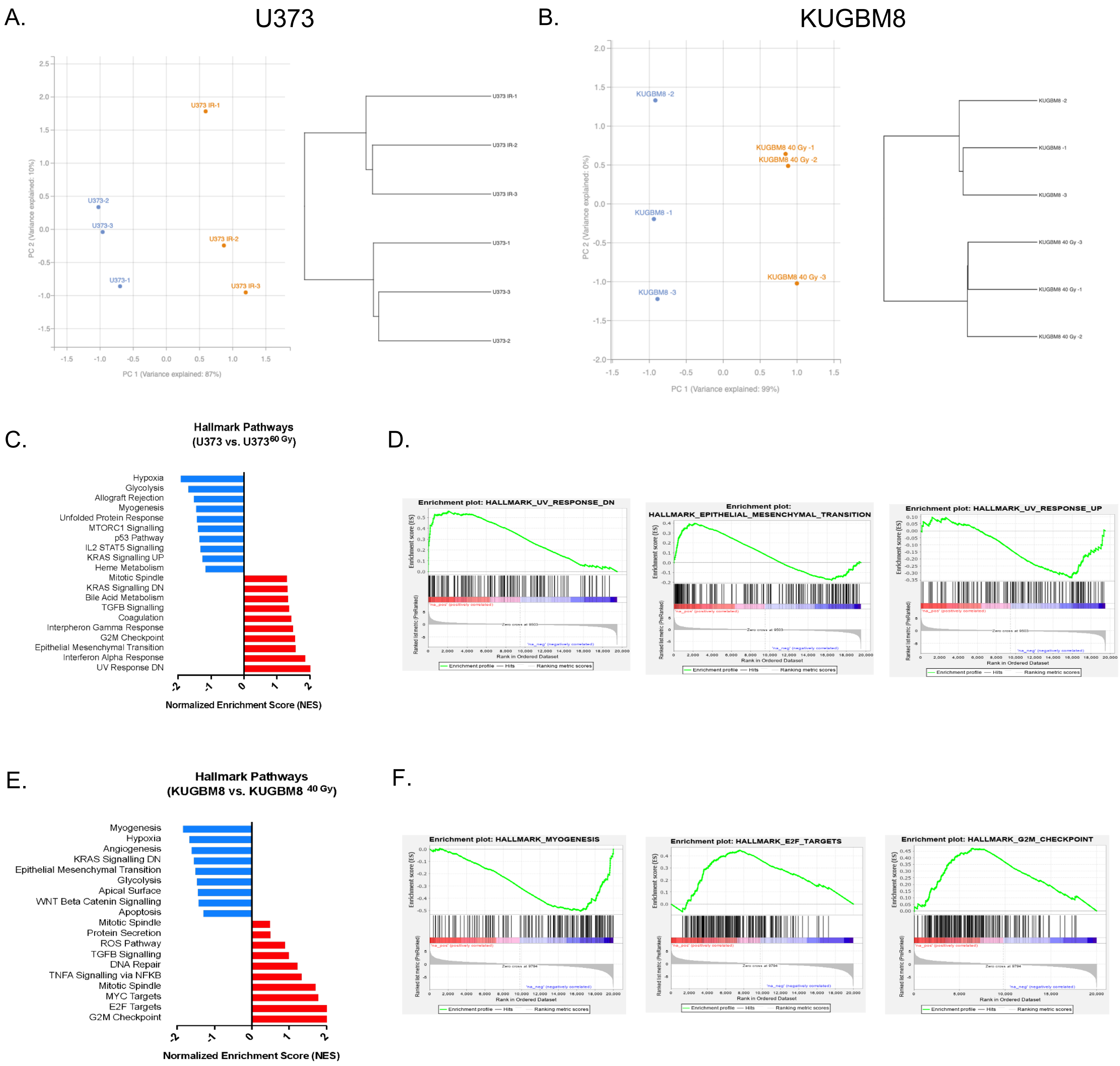


**Supplementary Figure S2.** A,B. Principal component analysis (PCA) plots and representations of hierarchical clustering result of U373-U373$^{60\ Gy}$ and KUGBM8-KUGBM8$^{40\ Gy}$ samples. C. Hallmark GSEA and representative enrichment plots of U373 cells. D. Hallmark GSEA and representative enrichment plots of KUGBM8 cells.

A.

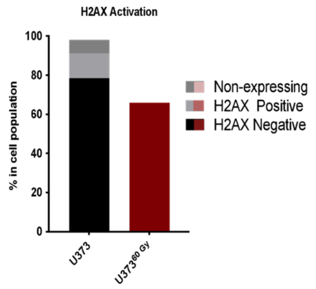


B.

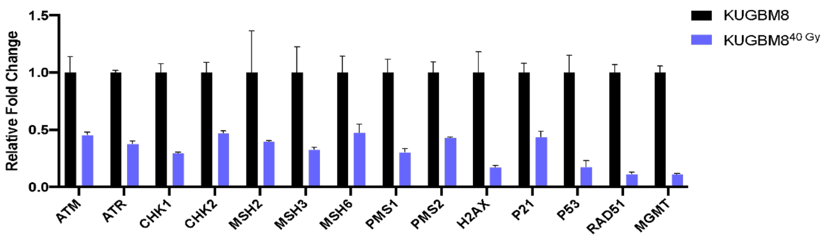


**Supplementary Figure S3**. A. Basal H2AX activation levels of U373 and U373$^{60\ Gy}$ cells. B. qRT-PCR results showing gene expression levels of different DNA damage response and repair elements in KUGBM8 cells.

| Gene | Sequence of Forward Primer (5’-3’) | Sequence of Reverse Primer (5’-3’) |
|---|---|---|
| GAPDH | AGCCACATCGCTCAGACAC | GCCCAATACGACCAAATCC |
| ATM | CATCGCATGTGATTAAAGCA | TTCTGATAGGAATCAGGGC |
| ATR | CAATTGTGGAGGAGATTTCC | CTTCTGAGAACTCTTGTATCTG |
| CHK1 | CGGTATAATAATCGTGAGCG | TTCCAAGGGTTGAGGTATGT |
| CHK2 | CTCGGGAGTCGGATGTTGAG | GAGTTTGGCATCGTGCTGGT |
| MSH2 | TTGATGGCCAGAGACAGGTT | CCATGTCTCCAGCAGTCTCT |
| MSH3 | CCATCATGGCTCAGATTGGC | ATCATGAGTGCTCGTCCCTC |
| MSH6 | ACTTCAGCAGGGACGTAACA | ACTTCAGCAGGGACGTAACA |
| PMS1 | ATGAGTGGAGCAGGGGAAAT | CACTTGCTGACATGGGTTTCT |
| PMS2 | ACTTCCGTGGATTCTGAGGG | GTGTTTGGGGTTGCGAGATT |
| MLH1 | CTACTTCCAGCAACCCCAGA | AGAACCTCATGTCCCTGCTC |
| H2AX | ACTCAACTGGGCAATCCAAG | GGGTTAGCTGCAGAATTCCA |
| p21 | GGCAGACCAGCATGACAGATTT | AAGATGTAGAGCGGGCCTTTGA |
| TP53 | GGTGACACGCTTCCCTGGATT | AGGGGGACAGAACGTTGTTTTCAG |
| RAD51 | TGCGGACCGAGTAATGGCA | TCCTTCTTTGGCGCATAGGCA |
| VEGFA | TGTGAATGCAGACCAAAGAAAGA | ACCAACGTACACGCTCCAG |
| MGMT | GGATTGCCTCTCATTGCTCC | CCCGTTTTCCAGCAAGAGTC |

**Supplementary Table S1.** List of primers and sequences used in qRT-PCR

| Antibody | Brand/Catalog Number |
|---|---|
| Phospho-ATR Ser428 | Cell Signaling/5161 |
| ATR Total | Cell Signaling/2790 |
| Phospho-ATM Ser1981 | Cell Signaling/13050 |
| ATM total | Cell Signalling/11G12 |
| Phospho-CHK1 Ser345 | Cell Signalling/133D3 |
| CHK1 | Cell Signalling/23605 |
| Phospho-CHK2 Ser19 | Cell Signalling/2666 |
| CHK2 | Cell Signalling/2662S |
| $\gamma$H2AX | EMD Millipore/05-636 |
| RAD51 | Abcam/ab213 |
| MSH2 | Abcam/ab52266 |
| MSH6 | Abcam/ab92471 |
| PMS1 | Abcam/ab129020 |
| PMS2 | Abcam/ab110638 |
| MLH1 | Abcam/ab92312 |
| MSH3 | Abcam/ab111107 |
| $\alpha$-tubulin | Abcam/ab15246 |
| GAPDH | Abcam/ab9485 |

**Supplementary Table S2.** List of primary antibodies used in Western Blot

| Antibody | Brand/ Catalog Number |
|---|---|
| Anti-vinculin | Abcam/ab73412 |
| Rhodamin-phalloidin | Thermo R415 |
| Mounting medium with DAPI | Abcam/ab104139 |
| Phosphor-Histone H2A.X (Ser139) | Millipore/05-636 |
| 53BP1 | Novus/NB100-304 |
| Alexa Fluor® 488 Anti- IgG Secondary Antibody | Invitrogen |
| Alexa Fluor® 594 Anti- IgG Secondary Antibody | Invitrogen |

**Supplementary Table S3.** List of primary and secondary antibodies used in immunofluorescence staining
